# Expansion of pneumococcal serotype 23F and 14 lineages with genotypic changes in capsule polysaccharide locus and virulence gene profiles post introduction of pneumococcal conjugate vaccine in Blantyre, Malawi

**DOI:** 10.1101/2024.03.25.586540

**Authors:** Rory Cave, Akuzike Kalizang’oma, Chrispin Chaguza, Thandie S. Mwalukomo, Arox Kamng’ona, Comfort Brown, Jacquline Msefula, Farouck Bonomali, Roseline Nyirenda, Todd D. Swarthout, Brenda Kwambana-Adams, Neil French, Robert S. Heyderman

## Abstract

Since the introduction of the 13-valent pneumococcal conjugate vaccine (PCV13) in Malawi in 2011, there has been persistent carriage of vaccine serotype (VT) *Streptococcus pneumoniae*, despite high vaccine coverage. To determine if there has been a genetic change within the VT capsule polysaccharide (cps) loci since the vaccine’s introduction, we compared 1,022 whole-genome-sequenced VT isolates from 1998 to 2019. We identified the clonal expansion of a multidrug-resistant, penicillin non-susceptible serotype 23F GPSC14-ST2059 lineage, a serotype 14 GPSC9-ST782 lineage and a novel serotype 14 sequence type GPSC9-ST18728 lineage. Serotype 23F GPSC14-ST2059 had an I253T mutation within the capsule oligosaccharide repeat unit polymerase Wzy protein, which is predicted *in silico* to alter the protein pocket cavity. Moreover, serotype 23F GPSC14-ST2059 had SNPs in the DNA binding sites for the cps transcriptional repressors CspR and SpxR. Serotype 14 GPSC9-ST782 harbour a non-truncated version of the large repetitive protein (Lrp), containing a Cna protein B-type domain which is also present in proteins associated with infection and colonisation. These emergent lineages also harboured genes associated with antibiotic resistance, and the promotion of colonisation and infection which were absent in other lineages of the same serotype. Together these data suggest that in addition to serotype replacement, modifications of the capsule locus associated with changes in virulence factor expression and antibiotic resistance may promote vaccine escape. In summary, the study highlights that the persistence of vaccine serotype carriage despite high vaccine coverage in Malawi may be partly caused by expansion of VT lineages post PCV13 rollout.

**Impact Statement:** Our findings highlight the potential for clonal expansion of multidrug-resistant, penicillin-non-susceptible vaccine serotype lineages with capsule locus modifications, within a high carriage and disease burden population. This shift has occurred among young children where there has been high vaccine coverage, posing challenges for effective vaccine scheduling and design. Furthermore, this study emphasises the importance of ongoing *Streptococcus pneumoniae* genomic surveillance as new or modified pneumococcal vaccines are implemented.

**2. Data summary:** Whole genome sequencing assemblies for the PCVPA survey have been deposited in the BioProject PRJNA1011974.

## 3. Introduction

Pneumonia, meningitis and sepsis caused by *Streptococcus pneumoniae* (the pneumococcus) is a major global public health concern, with an estimated 300,000 global deaths due to invasive disease reported among children aged under 5 of which 57% occur within resource poor settings (1,2). Asymptomatic nasopharyngeal carriage of the pneumococcus is a prerequisite for pneumococcal disease, transmission, and the development of natural immunity (3). Since the introduction of pneumococcal conjugate vaccines (PCVs) in childhood vaccination programs worldwide, there has been a considerable reduction in invasive disease (4).

The outer capsule polysaccharide (cps) is a major pneumococcal virulence factor, protecting the pneumococcus against desiccation, complement-mediated opsonophagocytic and other host antimicrobial pathways (5–7). The cps biosynthesis genes are found on a single locus controlled by a single promoter region for most serotypes (8). Cps consists of diverse sugar structures that vary among isolates, serving as the basis for classifying *S. pneumoniae* serotypes, with more than 100 immunologically-distinct serotypes identified to-date (9). Pneumococcal conjugate vaccines (PCVs) are formulated with a select array of serotype-specific capsule polysaccharides, chosen to target the most commonly occurring invasive serotypes, with a particular focus on those that cause the most severe diseases or are associated with AMR (10).

Since the 2011 PCV13 rollout in Blantyre, Malawi, we have shown a reduction in vaccine serotype (VT) invasive pneumococcal disease (IPD) with the incidence of post-PCV13 VT IPD 74% lower among children aged 1–4 years, and 79% lower among children aged 5–14 years from 2006 to 2018 (11). However, among PCV13 age ineligible populations, we noted only 38% lower VT IPD among infants and 47% lower among adolescents and adults. We have also shown that VT IPD has persisted amongst infants <90 (12). Alongside this, we have shown that in contrast to high income settings, there is considerable residual VT carriage, 7 years after PCV13 introduction despite high vaccine uptake (13). We have also shown that this imperfect direct and indirect control of pneumococcal carriage and disease is associated with waning of protective vaccine-induced anti-pneumococcal immunity in the first year of life (14).

The pneumococcus is highly transformable such that the *cps* locus, a known recombination hotspot, often acquires changes and can facilitate pneumococcal vaccine escape(15). Serotype switching from VT to non-vaccine serotypes (NVT), gene deletions or mutations resulting in pseudogenes lead to capsule loss (16,17). Genetic changes within the cps locus which do not lead to capsule switch or loss could also enable VT serotype persistence post PCV rollout (18). Here, we investigated the hypothesis that the residual VT IPD and persistent VT carriage observed following PCV13 introduction in Malawi, is at least in part due to the clonal expansion of VT lineages that have acquired changes in their *cps* locus while maintaining their serotype. We further postulate that these capsule locus variants have also acquired genetic traits that together could promote a competitive advantage in colonisation and transmission, with the potential to result in vaccine escape.

## 4. Methods

### Whole genome sequences

We obtained the whole genome sequences of PCV13 VT *S. pneumoniae* isolates from Blantyre, Malawi through the Global Pneumococcal Sequencing Project (GPS) dataset (n=329). These were derived from isolates collected between 1998-2015, including both carriage and disease isolates. We also used the Pneumococcal Conjugative Vaccine Prospective Analysis (PCVPA) dataset (n=693) from serologically typed carriage isolates collected from vaccinated, unvaccinated children, and adults living with HIV between 2015 to 2019 (13,19,20). Additionally, for the purpose of genomic comparison between Blantyre VT isolates with those isolated in other countries we incorporated publicly available isolates from Pathogenwatch which contains sequences from carriage and disease from early 1900’s to 2021 (https://pathogen.watch/, Accessed February 2023).

### Genome genetic typing and annotation

Genetic typing and antimicrobial resistance genotype of isolates was also conducted using Pathogenwatch. Lineages were defined by Pathogenwatch using the Global Pneumococcal Sequence Cluster (GPSC) nomenclature employing the PopPUNK framework, and the Multi-locus Sequence Type (MLST) system based on the pneumococcal scheme (19,21,22). Full genome annotation was conducted using Bakta v1.9.1 (23).

### *cps locus* comparison

We used parsnp v1.7.4 (https://github.com/marbl/parsnp) to extract and identify single nucleotide polymorphisms (SNPs) within the Blantyre pneumococcal *cps* locus by aligning them against reference serotype specific *cps* locus described by Bentley *et al*. (8). These SNPs were then annotated to distinguish synonymous from non-synonymous mutations using the vcf-annotator tool v0.5 (https://github.com/rpetit3/vcf-annotator). A phylogenetic tree from the alignment of the *cps* locus was constructed using IQ-TREE v2.1.2 with the best model for each alignment selected by ModelFinder (24,25). Phylogenetic trees and SNPS within the *cps* locus were visualised using the R package ggtree v3.18 (26). BLASTP v2.14.1 was used to determine if similar mutations were found in the amino acid sequence of isolates from other countries (27).

We performed sequence alignment for the *cps* locus using BLASTN v2.14.1 to enhance the detection of indels within the *cps* locus that are segmented into multiple contigs. The alignment coverage against the reference was then visualised using the R package gggenomes v0.9.12 (https://github.com/thackl/gggenomes).

To determine genetic synteny within the intergenic regions of the *cps locus*, we extracted the DNA sequences located between *dexB* and *wzg* for each individual genome. Synteny was established by aligning these sequences with one another using minimap2, and the resulting synteny patterns were visualised using the R package gggenomes (28). We investigated alterations within the 37-CE region located upstream of the cps promoter, where the transcriptional factors SpxR and CpsR are known to bind to suppress expression of the cps locus (29). To locate and extract the 37-CE sequence, we developed a custom Python script (https://github.com/rorycave/37-CE_finder) that identified the position of the sequence pattern “TTGAAAC,” which is typically conserved in the 37-CE region across various serotypes within the *cps* locus intergenic region. Subsequently, the 37-CE sequences were extracted using bedtools v2.28 ‘getfasta’ commands based on their sequence positions, including 156 nucleotides upstream and 14 nucleotides downstream of the sequence (30). The extracted sequences were then manually checked by aligning them to the reference 37-CE sequence using Clustal Omega v1.2.4 (29,31).

### The impact of synonymous SNPs on protein structure within the *cps* locus

To determine the impact of nonsynonymous SNPs on a protein structure within the *cps* locus, we first used SWISS-MODEL, to find homologous protein models that had a Global Model Quality Estimate (GMQE) >0.95 (32). Furthermore, within that GMQE range we chose X-ray diffraction models if present over models predicted by in silico methods. For transmembrane proteins that only have in silico predicted models, TMBed was used to identify if mutations occurred in the transmembrane cytoplasmic or extracellular domain of the protein (33). Additionally, we used PrankWeb 3 web server which runs P2RANK to determine if the mutation occurred in a protein pocket, and Missesne3D to assess whether mutations would cause structural damage to the protein’s 3D structure (34–36).

### Phylogenetic and accessory genome analysis

A core SNP maximum-likelihood (ML) phylogenetic tree was constructed by aligning assemblies to a serotype specific complete reference genome sequence using Snippy v4.6.0 (https://github.com/tseemann/snippy). Recombination within aligned sequences was then filtered out with Gubbins v3.3.1 (37). A phylogenetic tree was then constructed from recombinant-free alignment using IQ-TREE v2.1.2 with the best model for each alignment using selected by ModelFinder and set ultrafast bootstrap replication to 1,000 (24,25). The phylogenetic tree was visualised and annotated in Microreact (38).

To find differences in the accessory genome between lineages that have the same serotype, a pangenome form the annotated genome for each serotype was then constructed using Panaroo v1.3.4 with the merge paralogs setting (39). Scoary v1.6.16 was then used to identify genes that belong to certain lineages that were absent in others (40).

## 5. Results

### Expansion of cps locus variant lineages of serotype 23F and 14 following PCV13 introduction

To determine if the predominate VT cps locus genotypes changed after PCV13 introduction in Malawi, we compared the PCVPA dataset (2015 to 2019) to earlier pneumococcal sequences (1998 to 2014) in Blantyre, Malawi. Serotype 23F and 14 lineages were found to have expanded with changes in their *cps* locus genotypes.

The dominant serotype 23F lineage shifted from GPSC20 (n=23, 69.7%) pre-PCV13 rollout to a GPSC14 lineage that dominant from 2014 onwards (n=57, 78.03%). This includes the emergence of a GPSC14 ST2059 (n=42, 57.5%) that became dominant among all 23F serotype genotypes collected in the PCVPA dataset.

For serotype 14 isolates (n=77, collected between 2000 and 2019), the dominant GPSC9 remained unchanged after the PCV13 introduction (Figure 1B). However, there was a shift in the most prevalent STs, transitioning from GPSC9 ST63 pre-vaccine (n=13, 100%) to GPSC9 ST782 (n=20, 35.1% of PCVPA serotype 14 isolates) between 2015 and 2019. We also observed the clonal expansion of a novel serotype 14 sequence type ST18728 (n=13, 26.3% of PCVPA serotype 14 isolates), which was dominant from 2018 to 2019.

**Figure 1:**
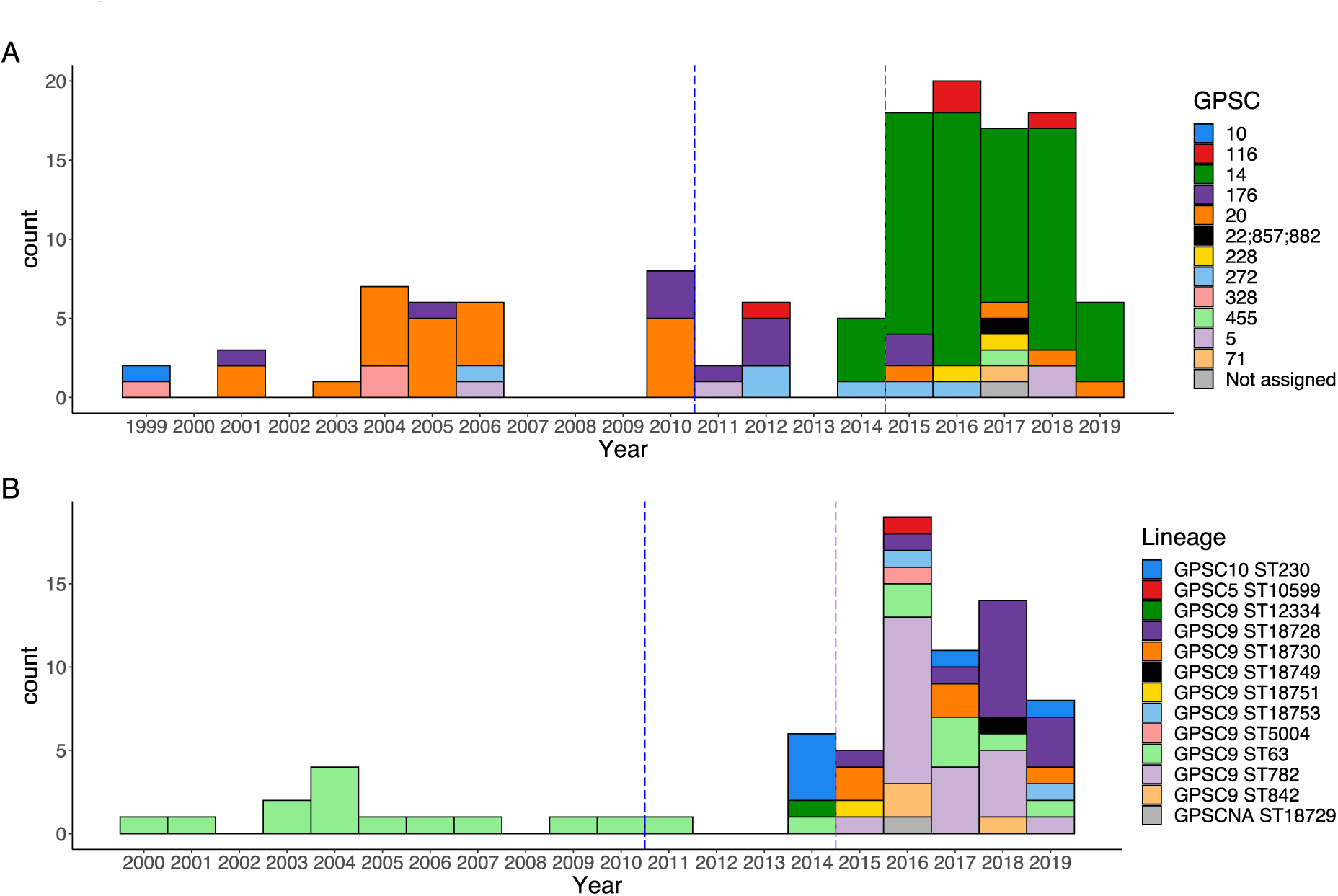
Change in genetic lineage/strain overtime among A) Serotype 23F and B) Serotype 14 isolates. Blue dash line indicates the time that PCV13 was introduced in Blantyre, Malawi. Purple line indicates the start of PCVPA survey.

### Genetic drift within 23F GPSC14 *cps* locus predicted to impact on capsule expression and phenotype

To explore the hypothesis that the clonal expansion of 23F GPSC14 ST2059 post-PCV13 introduction and the decline in 23F GPSC20 may have been due a competitive advantage, we first looked for genetic differences within the *cps* locus. Alignment with the reference 23F *cps* locus (GenBank accession: CR931685) revealed a higher SNP density with the emergent 23F GPSC14 lineage (mean = 11.5 SNPs/Kbp, range 7.7 - 29.2 SNP/Kbps per isolate) when compared to 23F GPSC20 (mean = 5.7 SNPs/Kbp, range 5.7 to 5.8 SNPs/Kbp per isolate). Furthermore, the 23F GPSC14 lineages exhibited four non-synonymous SNPs, resulting in amino acid changes: W54S, L56F, T237I, and I253T, in the oligosaccharide repeat unit polymerase gene, *wzy*. These same mutations were also found in a single serotype 23F GPSC455 isolate (Figure 2).

**Figure 2:**
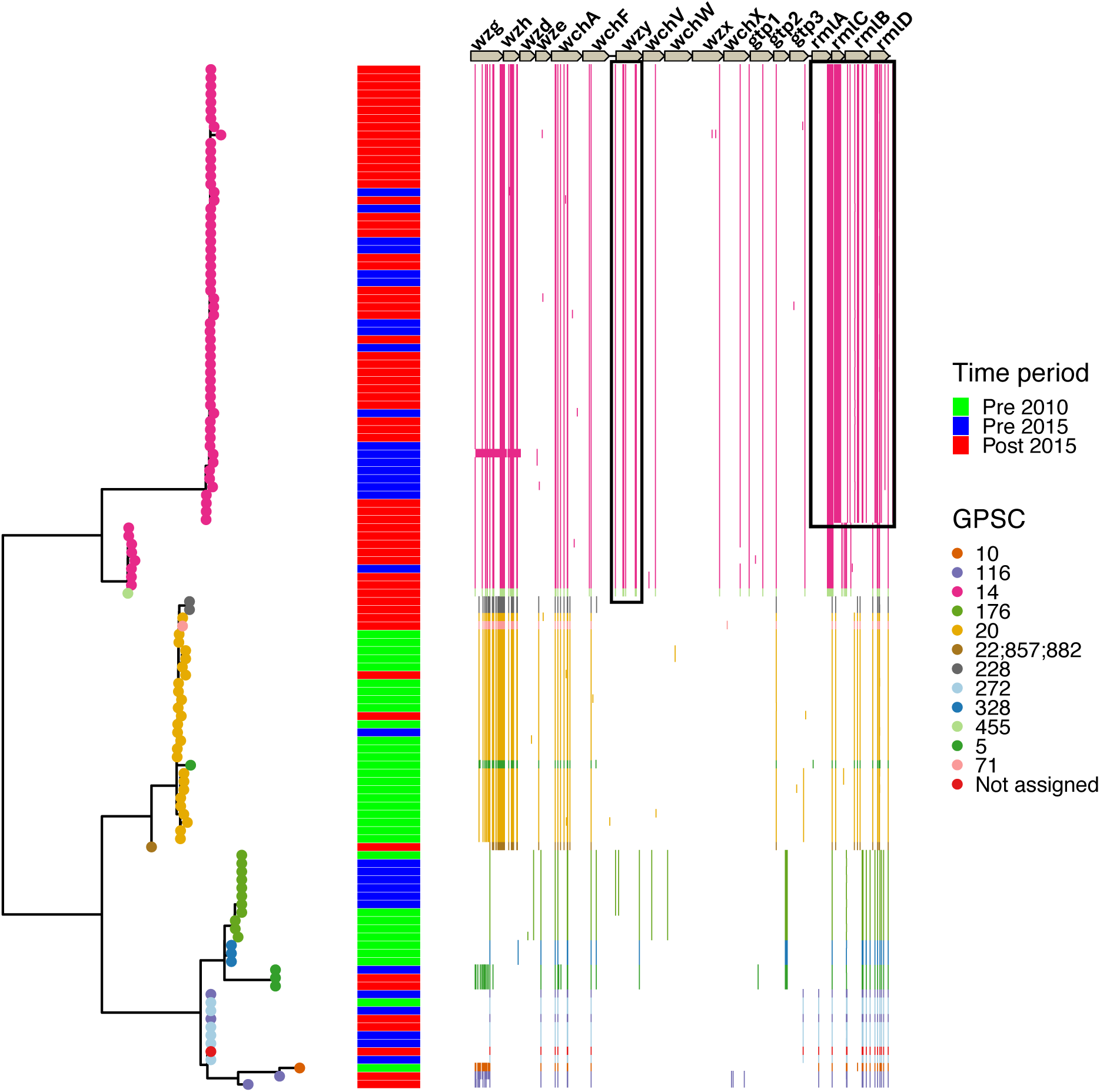
Maximum likelihood phylogenetic tree and position of SNPS within serotype 23F *cps* locus shows that change in genotype in *wzy* and *rmlACBD* in the emergent GPSC14 *S. pneumoniae* isolates in Blantyre, Malawi based on phenotype clustering.

Additionally, from the cps locus alignment, we identified a subpopulation within the GPSC14 lineage composed of ST2059 (n=49), ST12347 (n=1), and isolates (n=7) that belong to five novel sequence types (ST18717, n=1; ST18707, n=2; ST18727, n=1; ST18722, n=1; ST18723, n=2). These were single allele variants of ST2059 that displayed recombination within the rhamnose synthesis locus (*rmlACDB*). This resulted in three non-synonymous SNPs (leading to amino acid changes: L214V, R258M, and S272P) in the *rmlA* gene and seven non-synonymous SNPs (leading to amino acid changes: T2S, A12V, E13I, L46E, E57A, D89G, and STOP198E) in the *rmlC* gene.

To evaluate the impact of non-synonymous SNPs in 23F GPSC14 on protein structure, we utilised an AlphaFold model for *S. pneumoniae* Serotype 23F Wzy (UniProt ID: Q9R925), an Alphafold model of RmlA (UniProt ID: A0A2I1UG67) from *Streptococcus oralis subsp dentisani* and a X-ray diffraction model for RmlC (SMTL ID: 1ker.1.) of *Streptococcus suis* (32). From these three, the I253T mutation in Wzy was predicted to be part of a protein pocket (confidence score of 0.970), and the alteration of the amino acid residue was predicted to induce structural changes in the protein by reducing the cavity volume by 126.576 Å^3^ (Figure S1).

We then investigated the intergenic regions of the serotype 23F *cps* locus, situated between *dexA* and the *wzg* gene where transcriptional regulators bind to alter capsule expression (27). We observed structural variations among several different lineages (Figure 3A). GPSC14 and GPSC455 exhibited a longer intergenic region (1,401bp) compared to GPSC20, 228, 5, and 22 (554bp) due to the truncation of insertion sequences. Conversely, GPSC176, 272, 328, 116, 22, and 40 displayed a shift in the repeat unit pneumococcal (RUP) preceding the insertion sequences. These structural variations in the intergenic regions of the cps locus may lead to different level of gene expression in the cps locus.

**Figure 3:**
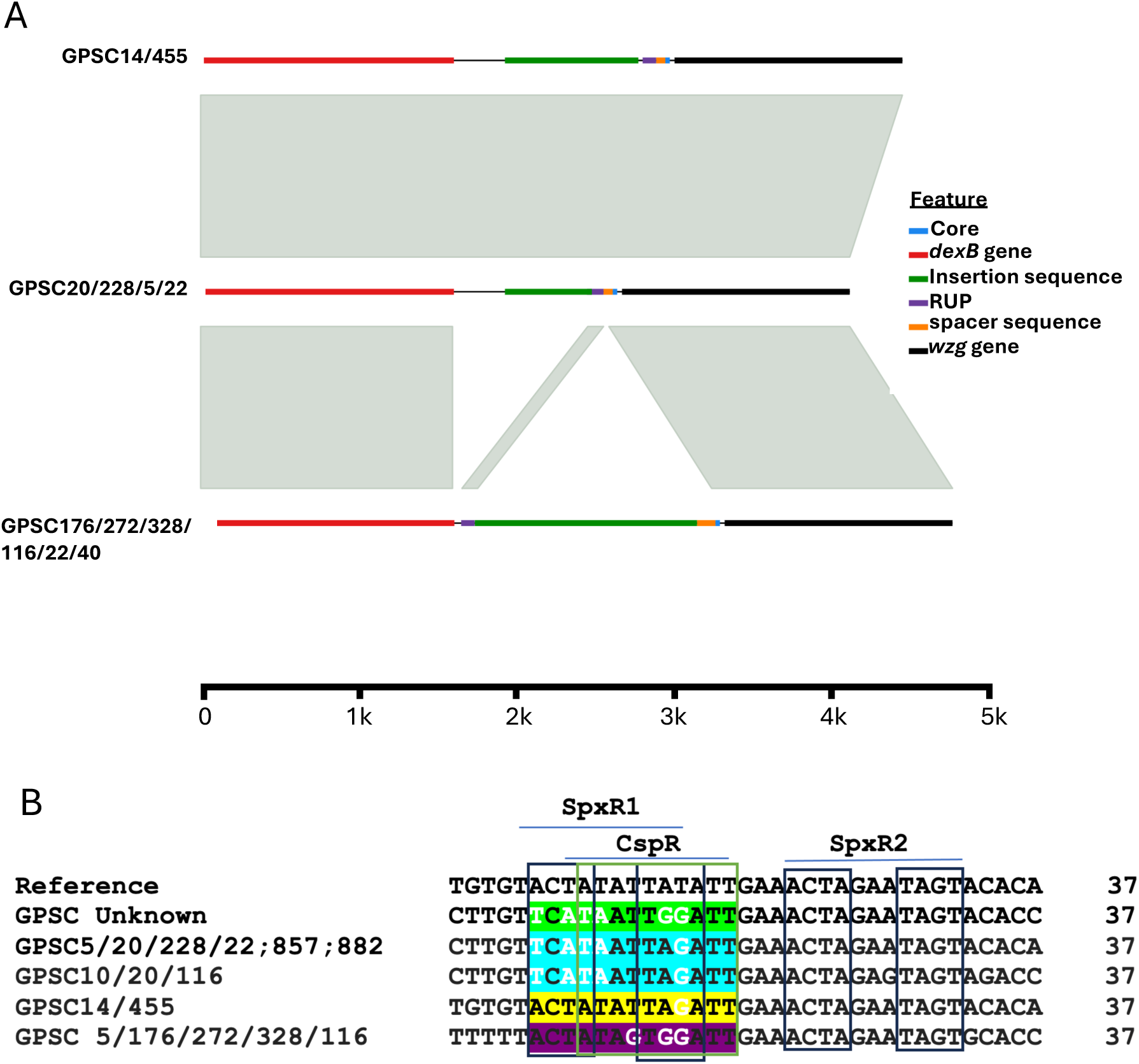
Differences in the *cps* locus intergenic region in the emergent 23F GPSC14 isolates in Blantyre, Malawi. A) Genetic synteny of intergenic regions between isolates. B) Changes in the CpsR and SpxR1 nucleotide binding site in the 37-CE.

Additionally, we identified differences between lineages in DNA binding site sequences for SpxR and CpsR in the 37-CE, which are known to suppress capsule expression (8) (figure 3B). Notably, GPSC14 and GPSC455 showed a higher similarity (92%) to the reference strain (D39 Serotype 4) in the SpxR1 and CpsR binding sites described by Glanvielle *et al.*, compared to GPSC20, 228, 22, 857, 882, 10, and 116 (62% similarity). Moreover, within GPSC5, 176, 272, 328, and 116 there was 72% similarity compared to the reference sequence. The primary distinction between the GPSC20 SpxR1 and CpsR binding sites and the reference sequence lies in the alterations involving A to T or T to A base changes at positions 1, 3, 4, and 5, along with a T-to-G substitution at position 10. Conversely, for GPSC14 and GPSC455, the sole variation from the reference occurs at position 10, where a T is replaced by a G, whereas GPSC5, 176, 272, 328, and 116 also had two additional changes at positions 7 and 9, both being A to G. Ultimately these changes may alter the binding affinity of the SpxR1 and CpsR proteins, leading to changes in cps locus gene expression.

### Emergent serotype 14 isolates with the non-truncated version of the large repetitive protein gene within the *cps* locus

Through a comparative analysis of the serotype 14 *cps* locus over time, we did not observe the emergence of non-synonymous SNPs within cps genes or changes in cps transcriptional binding sites, that we had seen in the 23F lineage. Instead, by BLAST sequence analysis using a reference serotype 14 sequences (GenBank accession: CR931662) we identified isolates harbouring large repetitive protein (*lrp*) gene that was not truncated, containing the Cna protein B-type domain a feature absent from serotype 14 isolates *lrp* gene collected before 2015 (figure 4). The emergence of isolates with the complete version of the *lrp* gene post-PCV13 introduction may suggest that the bacteria have a fitness advantage, potentially aiding them in evading vaccine-induced antibodies through alterations in the bacteria’s immunogenicity.

**Figure 4:**
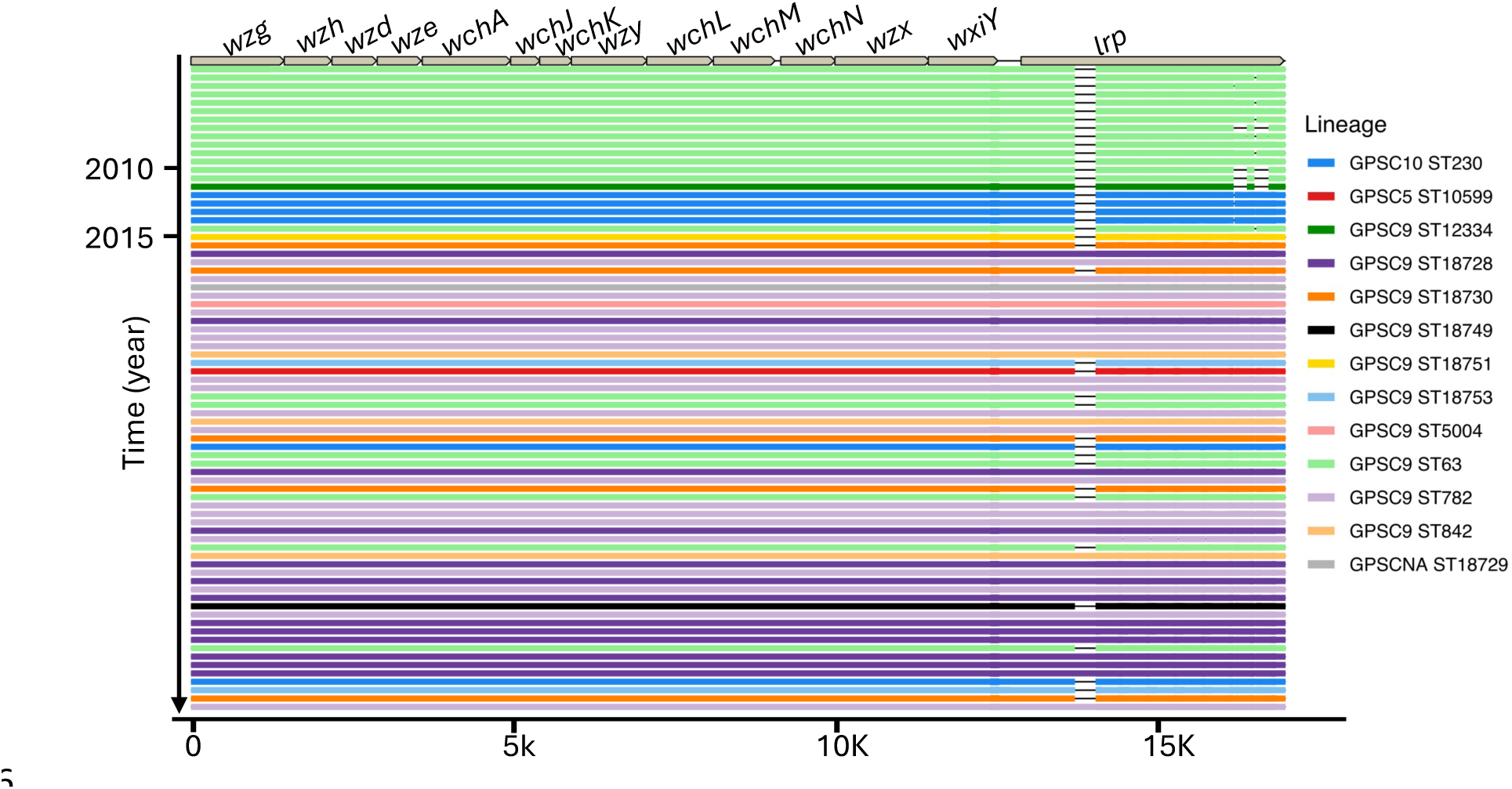
BLASTN analysis to identify large genomic changes in the serotype 14 cps sequences showing the emergence of sequence types within GPSC9 lineage that have complete version of *lrp* gene. GPSCNA= non-classified GPSC.

### Global comparative analysis of serotype 23F Wzy, SpxR1 and CpsR binding motif, and serotype 14 Lrp changes

To further understand whether the mutations related to the serotype 23F Wzy amino acid structure and the SpxR1 and CpsR binding sites in 37-CE are specific to certain lineages, and if they are confined to Malawi, we conducted genomic comparisons with 1,385 isolates worldwide. Notably, mutations in Wzy protein sequences and the genetic structures of the 37-CE were consistent across the majority of GPSC14 isolates globally (Figure S2). Moreover, the Wzy protein in serotype 23F exhibits a high level of sequence conservation, with just two distinct versions of the protein found in more than one isolate within the available dataset. This suggests, that emergent of a mutated Wzy protein in GPSC14 isolates may confer a fitness benefit.

In our core maximum likelihood phylogenetic analysis of serotype 23F isolates, constructed by aligning against the complete genome of serotype 23F *S. pneumoniae* ATCC 700669 (GenBank accession: FM211187), we found that the 37-CE sequences are primarily associated with specific lineages (Figure S2). This implies that different lineages may be linked to varying levels of cps expression. The phylogenetic tree also suggests a close genetic relationship between the newly emergent GPSC14 ST2059 in Blantyre and GPSC14 ST2059 isolates from South Africa, with the closest South African Malawian isolate differing by only 32 SNPs. This implies possible country-to-country transmission, most likely from South Africa to Malawi, as the ST2059 lineage was prominent in South Africa pre and post-PCV vaccine introduction but not in Malawi.

We also conducted a comparative analysis of serotype 14 isolates’ *lrp* gene. Utilising data from 1,607 isolates collected globally, including Malawian isolates from various regions, our aim was to determine whether the non-truncated version of the *lrp* gene is associated with specific lineages. Interestingly, we observed that, aside from GPSC9, other lineages mainly had the complete or truncated version of the gene, indicating that the versions of the gene evolved separately from each other, and recombination is less likely to occur in the *lrp* gene.(table S1). Our phylogenetic analysis of GPSC9 serotype 14 isolates, aligned against the serotype 14 *S. pneumoniae* G54 complete genome (GenBank accession: CP001015), indicates a genetic divergence based on the completeness of the *lrp* gene. We observe one cluster containing all truncated versions of the *lrp* gene, and the other comprises a mixture of complete and incomplete *lrp* genes, suggesting that variations in the *lrp* gene could potentially play a role in the differences observed among these isolates (Figure S4).

### Emergent serotype 23F and 14 lineages are associated with virulence genes and AMR

To explore the hypothesis that the emerging serotype 23F and 14 lineages had additional virulence and AMR characteristics that conveyed advantage, we first compared the accessory gene content of the different lineages (20) (Table S2). A toxin-antitoxin system gene associated bacterial stress response, cell growth, and biofilm formation, and the pneumococcal serine-rich repeat protein (PsrP)-accessory Sec system (secY2A2) pathogenicity island witch involved in biofilm formation, adhesion to epithelial cells, and the export of contact-dependent pneumolysin toxin were consistently present in the Serotype 23F GPSC14 isolates but absent in the 23F GPSC20 isolates (41–47). Furthermore, all 23F ST2059 isolates carried the macrolide resistance gene *mefA*, which was only present in three 23F isolates outside of GPSC14. Additionally, they harboured Thiazolylpeptide-type bacteriocin and Lantibiotic resistance genes associated with bacterial colonisation (48).

Regarding serotype 14 GPSC9 lineage, the Choline-binding protein *pcpA* virulence gene which mediates pneumococcal adhesion was present in all but one isolate in cluster C, which contains the post-PCV13 emergent lineage on the core phylogenetic tree (Figure 5)(49). ST18728 also carried the Zinc metalloprotease virulence gene associated with inflammation in the lower respiratory tract, found in only one additional isolate indicating the emergent isolates may have an increased virulence potential enabling the lineage to the dominate lineages pre-PCV13 introduction (50).

**Figure 5:**
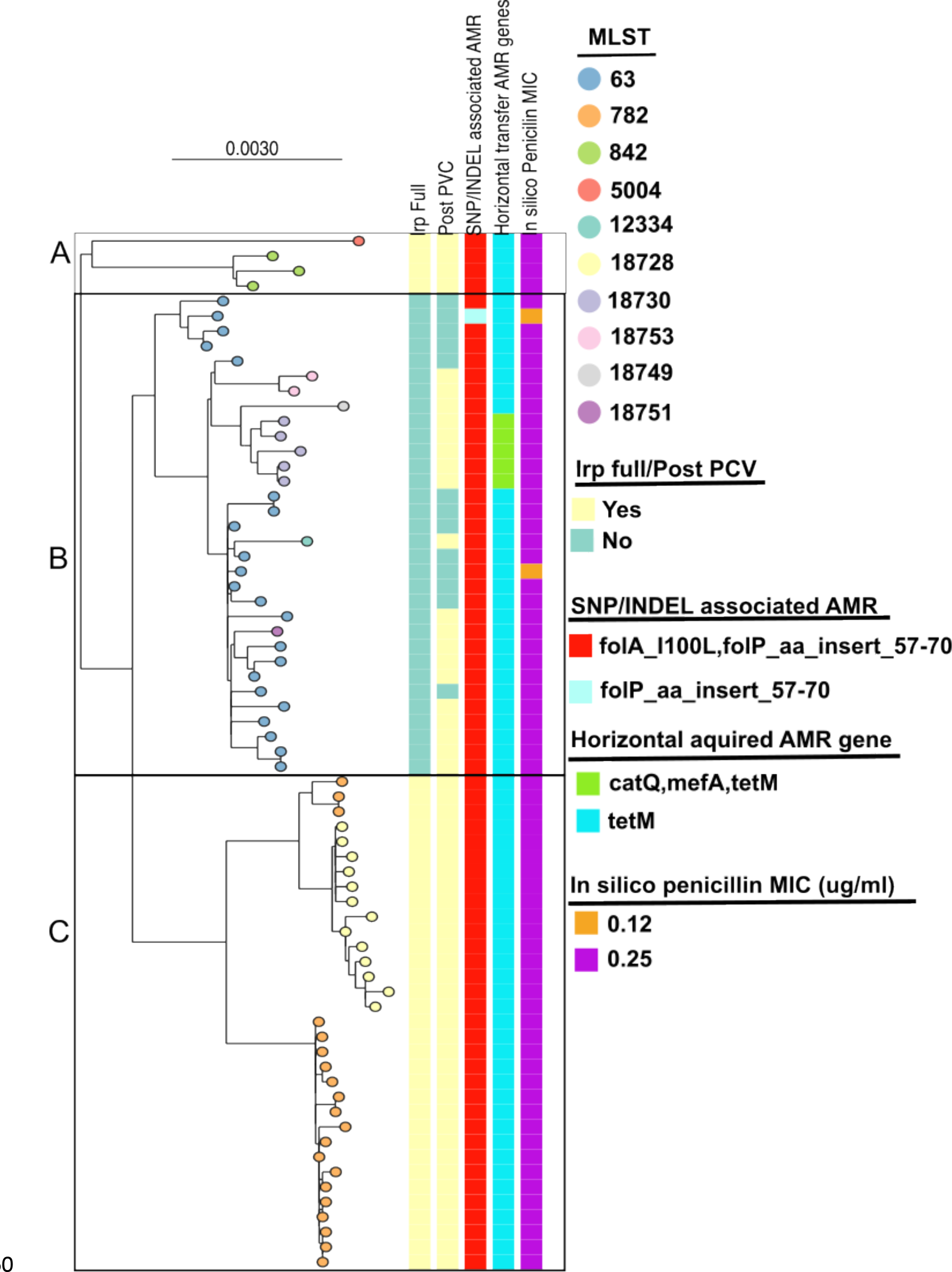
Phylogenetic tree of serotype 14 GPSC9 isolates from Blantyre, Malawi, and the clustering used to define different sub-lineages of GPSC9 for accessory gene content comparison.

To assess AMR among the emergent serotype 23F and 14 lineages, we compared them to the lineage that dominated before PCV 13 introduction (Figure 6). For 23F ST2059 harboured *mefA* (macrolide resistance), *tetM* (tetracycline resistance) and the mutations folP_aa_insert_57-70 and folA_I100L (resistance to sulfamethoxazole and trimethoprim, respectively). Additionally, 23F ST2059 isolates were penicillin non-susceptible (MIC 0.25 to 0.5 µg/µL), unlike GPSC20 ST82, which was penicillin-susceptible (MIC 0.035 µg/µL) (Figure 5). There were three 23F GPSC20 ST9530 isolates recovered before PCV13’s introduction also exhibited penicillin non-susceptibility (MIC 0.25 µg/µL).

**Figure 6:**
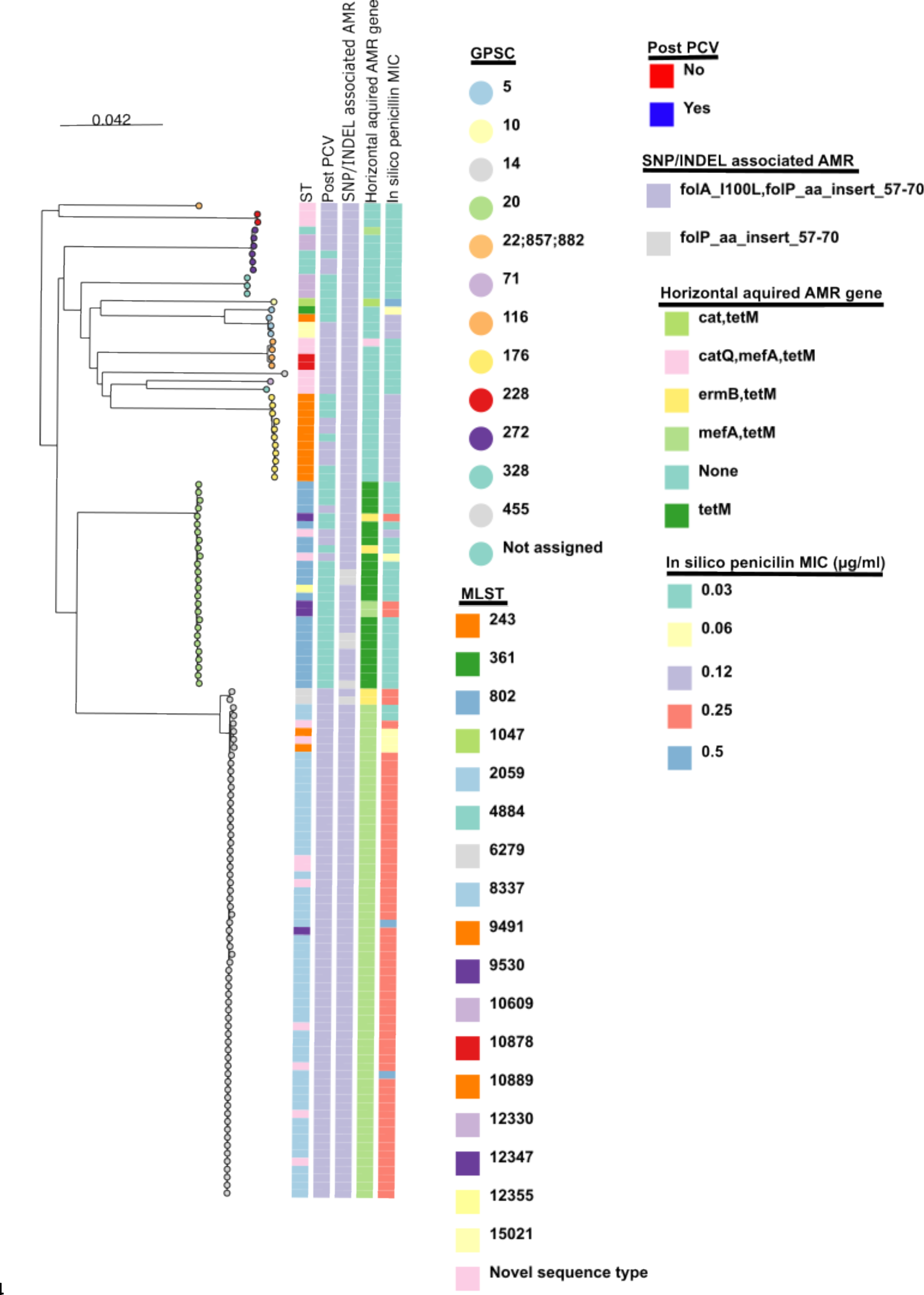
Phylogenetic tree of serotype 23F isolates from Blantyre, Malawi showing the emergent GPSC14 ST2059 isolates having resistance to more antibiotic classes from genotype data over other lineages.

For Serotype 14 isolates there were no additional gene mutations (folP_aa_insert_57-70 and folA_I100L) or acquisitions (*tetM*) conferring AMR. All serotype 14 lineages were penicillin non-susceptible. There was no additional increase in penicillin MIC amongst the emerging GPSC9 lineage.

## 7. Discussion

In the context of persistent *S. pneumoniae* VT carriage and high PCV13 uptake in Malawi, we have identified serotype 23F and 14 lineages, serotypes that have been prominent causes of IPD, with potentially functionally important genetic changes in their *cps* locus (13,51). These lineages harbour additional virulence attributes and AMR, which may further enhance their potential for vaccine escape. Indeed, together with a recently described serotype 3 GPSC ST700–GPSC10, clonal expansion of these lineages may explain the residual circulation of some VT serotypes after vaccination in Blantyre, Malawi (13). The detection of these lineages was only possible through continuous genomic surveillance at the population level, highlighting the importance of such surveys following PCV introduction.

The predominant serotype 23F lineage before PCV13 introduction in Malawi was GPSC20 ST802, a lineage found on multiple continents (52,53). However, post-PCV13 introduction, there was a decrease in ST802, and an expansion of GPSC14 ST2059, which, to date, has only been reported from neighbouring South Africa and Mozambique (52,54). Intriguingly, data from South Africa suggests that 23F ST2059 has a more invasive phenotype (19). Whether vaccine pressure was directly responsible for this shift is uncertain, as genotype replacement of 23F isolates has been reported in China (GPSC24 ST342 to multidrug-resistant GPSC16 ST81 also known as the PMEN1 clone lineage) which does not have a routine PCV programme (53). However, once established in Malawi, the genotypic attributes of the 23F GPSC14 ST2059 lineage may explain its persistence, particularly in the context of vaccine-induced serotype-specific immunity that has waned below the correlates of protection for both carriage and disease in the first year of life (14).

We have identified two mutations within the *cps locus* of the emergent 23F GPSC14 lineage that were absent in other 23F genetic lineages. These are predicted to have an impact on capsule expression and production(29,55). These mutations included Wzy, an oligosaccharide repeat unit polymerase responsible for linking sugar chains outside the cell wall as part of the capsule. Wzy shows significant diversity between serotypes but is highly conserved within serotypes (18,56). It was therefore unexpected to find that the 23F GPSC14 lineage has multiple non-synonymous SNPs, including a SNP I253T that reduces the cavity volume found within a protein pocket of Wzy. Mutagenesis studies of Wzy involved in lipopolysaccharide synthesis in gram-negative bacteria such as *Pseudomonas aeruginosa* and *Shigella flexneri*, affect the O-antigen chain length and distribution affect capsule thickness (57,58). Moreover, differences in the intergenic region structure, especially the SNP’s in the SpxR1 and CpsR DNA binding sites in the 37-CE have previously been shown to alter the level of capsule production affecting the strain’s virulence (29,55). Furthermore, recent phenotypic studies have revealed a 6.6-fold variation in cps production among 23F isolates, though the lineage and *cps locus* genotype was not explored (59). We therefore hypothesise that mutations in Wzy and 37-CE will most likely affect capsule function and expression respectfully, conferring a competitive advantage.

The prevalent serotype 14 lineage before PCV13 introduction in Malawi was GPSC9 ST63, a globally distributed sequence type (60). Post-PCV introduction, *S. pneumoniae* GPSC9 isolates from the USA, Israel, South Africa, and Cambodia have been identified that have switched serotype from 14 to nonvaccine type 15A (60). This has not been seen in Malawi, instead GPSC9 ST63 has been overtaken by an expanding serotype 14 GPSC9 ST782, another global sequence type (61,62). Moreover, there was also the expansion of phylogenetically closely related novel sequence type, ST18728, which to date has only been reported in Malawi. This expanding serotype 14 GPSC9 are associated with genetic alteration in the *lrp* gene, where we identified the emergent strain GPSC9 ST782 and a novel sequence type closely related to Blantyre’s ST782, which harbours the complete 1359-amino acid-long version of the Lrp protein containing the Cna protein B-type collagen-binding domain. Variants in the lrp gene have been previously reported, particularly the serotype 14-like isolates from Papua New Guinea (63,64). However, the Papua New Guinea isolates differed from Blantyre isolates, not reacting to serotype 14 antibodies, and lacking key genes. The Lrp protein’s function is unknown, but it’s hypothesised to be a dominant antigen, overriding serological similarities between serotype 14 and 15 (8). The Lrp protein’s collagen-binding Cna B domain, also found in the virulent factor RrgB protein, could affect serotype 14’s colonisation ability and immunogenicity (59,60).

In addition to changes in the *cps* locus, possible factors contributing to the emergence and spread of these 23F and 14 lineages include genes associated with AMR, colonisation, and virulence that would provide bacteria with an advantage over lineages from the same setting. The question is whether these lineages that seem to have a competitive advantage will disseminate more widely. It is noteworthy that a different serotype 23F lineage, PMEN1, identified before pneumococcal vaccines were widely introduced, became transmitted across the globe, causing invasive disease (65–67). The emergence and spread of PMEN1 were attributed to the acquisition of antibiotic resistance, transmission, and virulence genes absent in closely related ancestor strains (68). Together this shows the importance of the identifying emergent strains with genetic adaption and assessing their potential to transmit locally and globally.

The main limitation of this study is the limited number of genomes from Malawi prior to and immediately after the introduction of PCV13 and that these historical genomes were biased towards invasive rather than carriage isolates. However, we speculate that, based on closely related isolates from neighbouring countries, such as the ST2059 isolates in South Africa, that these expanding lineages are likely to cause invasive disease in Malawi (69,70).

In conclusion, following the introduction of PCV13 in Malawi, there has been clonal expansion of 23F and 14 lineages, characterised by genotypic *cps* locus changes that affect capsule expression and production, and potentially interaction with host immune system. These lineages also have genetic features that confer a competitive advantage in terms of colonisation, transmission and AMR. These findings underscore the value of robust *S. pneumoniae* genomic surveillance to inform vaccine programmes in high burden settings where, in the face of a high force of infection, sufficient control of pneumococcal colonisation and disease has not yet been achieved (11,71,72).

## Author statements

### 8. Author contributions

*[Mandatory for Access Microbiology, encouraged for all other journals. A section describing each author’s contribution to the research, using the CRediT taxonomy from CASRAI:* https://casrai.org/credit/]

### 11. Ethical approval

The original study protocol for the PCVPA dataset *was* proved by the College of Medicine Research and Ethics Committee, University of Malawi (P.02/15/1677) and the Liverpool School of Tropical Medicine Research Ethics Committee (14.056). Written informed consent was obtained from adult participants and parents/guardians of child participants. Additionally, children aged 8–10 years provided informed assent. The consent encompassed permission for publication.

## 12. Acknowledgements

[This section is optional for all journals. However, if materials and results were obtained from outside the authors’ laboratories (e.g. production of antibodies, properties of strains), this must be acknowledged.] This section should also be used to list the individuals who are part of a listed Group Author/Consortium].

