## Supplemental figures and tables for "Expansion of pneumococcal serotype 23F and 14 lineages with genotypic changes in capsule polysaccharide locus and virulence gene profiles post introduction of pneumococcal conjugate vaccine in Blantyre, Malawi"

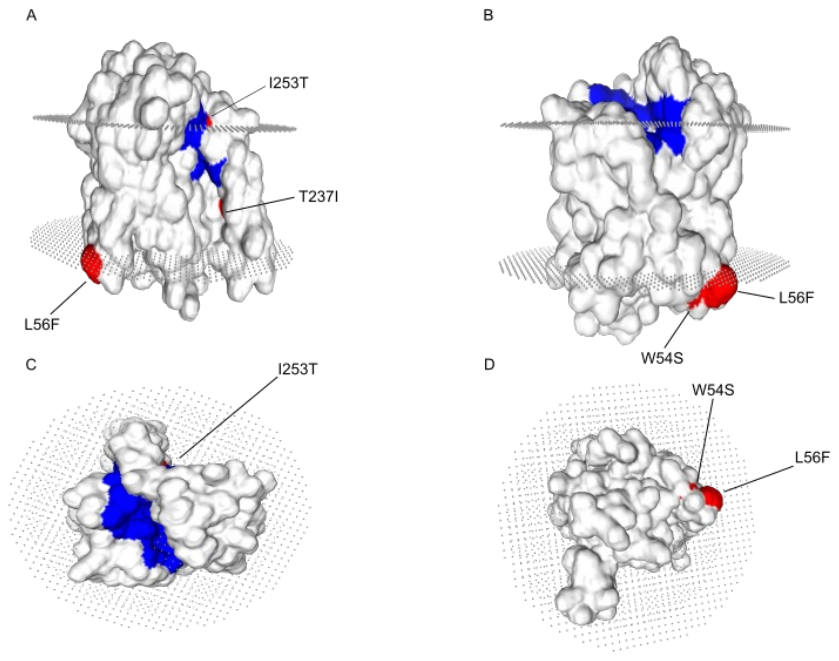
Figure S1: 3D Structure of Serotype 23F Wzy protein from an AlphaFold Model (PDB accession: Q9R925P), showing the I253T mutation which causes a structural change in the protein, found within a protein pocket. A) Front view B) back View C) Extracellular view D) Cytoplasmic view. Grey dots represent the bacterial membrane. Blue-highlighted amino acid residues represent the predicted protein cavity pocket, while red-highlighted amino acid residues represent amino acid changes from non-synonymous mutation.

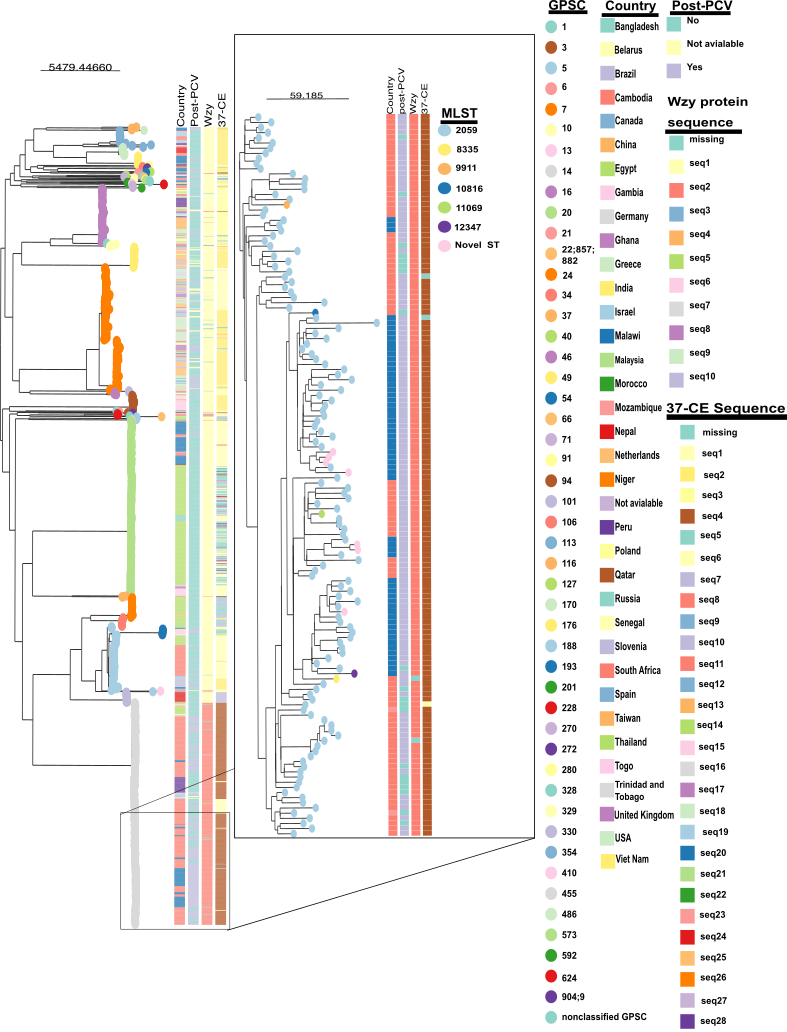

Figure S2: Serotype 23F core SNP Phylogenetic tree showing difference of Wzy protein sequences and 37-CE sequences in isolates from different lineages. Zoom in sections of the tree show phylogenetic close relationship of South African ST2059 to Malawian ST2059. Wzy protein sequence and 37-CE sequence can be seen in supplement figure S1 and S2 respectively.

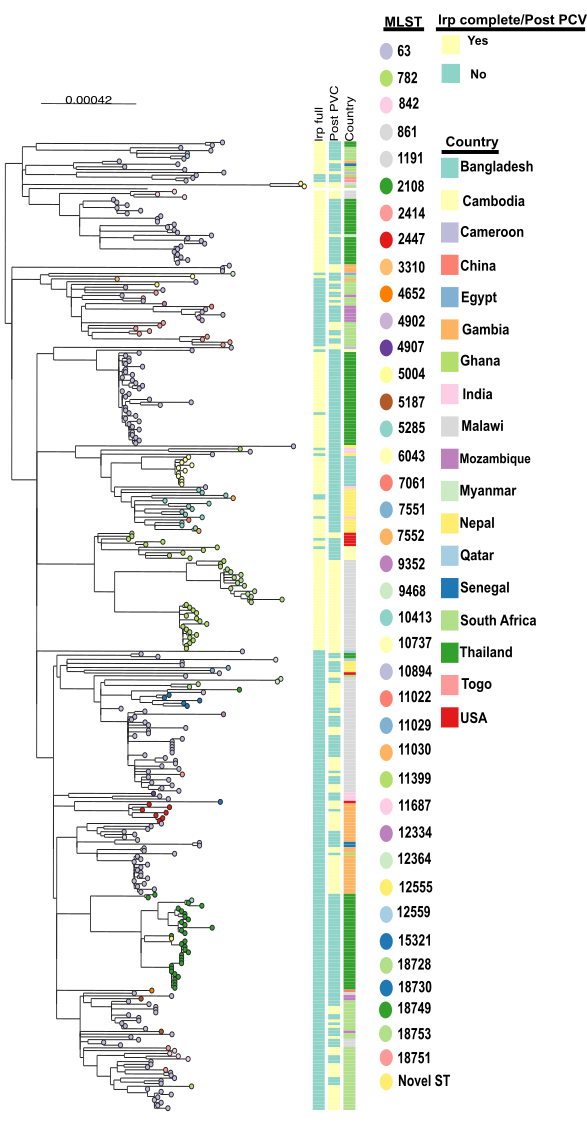

Figure S3: Serotype 14 GPSC9 core SNP phylogenetic tree showing the divergence in lineage related to have complete or truncated version of the *lrp* gene.

**Table S1: Distribution of complete and truncated versions of serotype 14 *lrp* gene across genetic lineages**

|  | Complete *lrp* gene | |
| --- | --- | --- |
| GPSC | Yes(%) | No (%) |
| 18 | 489 (98.39) | 8 (1.61) |
| 39 | 159 (100) | 0 (0) |
| 904;9 | 149 (40.93) | 215 (59.07) |
| Not assigned | 2 (66.67) | 1 (33.34) |
| 5 | 0 (0) | 1 (100) |
| 10 | 1 (0.62) | 158 (99.37) |
| 6 | 345 (99.71) | 1 (0.28) |
| 16 | 16 (100) | 0 (0) |
| 288 | 5 (100) | 0 (0) |
| 108 | 28 (96.55) | 1 (3.45) |
| 279 | 5 (100) | 0 (0) |
| 14 | 1 (50) | 1 (50) |
| 4 | 7 (100) | 0 (0) |
| 1 | 1 (100) | 0 (0) |
| 90 | 0 (1) | 1 (100) |
| 301 | 0 (0) | 4 (100) |
| 238 | 4 (100) | 0 (0) |
| 571 | 1 (100) | 0 (0) |
| 103 | 1 (100) | 0 (0) |
| 28 | 0 (0) | 71 (100) |
| 79 | 1 (100) | 0 (0) |
| 21 | 0 (0) | 1 (100) |
| 27 | 1 (100) | 0 (0) |
| 11 | 1 (100) | 0 (0) |

**Table S2: Selected accessory virulence genes present in the emergent serotype 23F and 14 lineages**

| Lineage/Strain | Gene function | Sensitivity | Specificity |
| --- | --- | --- | --- |
| 23F GPSC14 | Toxin-antitoxin system toxin component domain protein; CI-like repressor metallo-proteinase motif protein;Imm40 domain-containing protein; ImmA/IrrE family metallo-endopeptidase | 100.0 | 93.4 |
|  | accessory Sec system protein Asp2 | 100.0 | 78.7 |
|  | accessory Sec system protein translocase subunit SecY2 | 100.0 | 78.7 |
|  | accessory Sec system protein Asp1 | 100.0 | 78.7 |
|  | accessory Sec system protein Asp2 | 100.0 | 78.7 |
|  | accessory Sec system protein translocase subunit SecY2 | 100.0 | 78.7 |
|  | Glycosyltransferase GlyF; Glycosyl transferase family 8 | 100.0 | 78.7 |
|  | Glycosyl transferase family 2; Glycosyltransferase GlyG | 100.0 | 78.7 |
|  | accessory Sec system protein Asp3 | 100.0 | 78.7 |
|  | Glycosyltransferase GlyE | 100.0 | 78.7 |
|  | sugar transferase gtf3 | 100.0 | 78.7 |
|  | accessory Sec system translocase SecA2 | 100.0 | 77.0 |
|  | accessory Sec system glycosylation chaperone GtfB | 100.0 | 77.0 |
|  | accessory Sec system glycosyltransferase GtfA | 100.0 | 77.0 |
| 23F GPSC14 ST2059 | macrolide efflux MFS transporter Mef(A) | 100.0 | 76.6 |
|  | ABC-F type ribosomal protection protein Msr(D) | 100.0 | 76.6 |
|  | Lantibiotic biosynthesis protein | 100.0 | 72.7 |
|  | Lantibiotic efflux protein | 100.0 | 72.7 |
|  | Lantibiotic biosynthesis protein | 100.0 | 72.7 |
|  | Thiazolylpeptide-type bacteriocin | 100.0 | 72.7 |
| 14 GPSC9 cluster C | Choline binding protein PcpA | 97.0 | 97.3 |
| 14 GPSC ST18728 | Zinc metalloprotease ZmpB | 100.0 | 98.2 |

Sensitivity is the presence of this gene within target genotype and specificity is the absent of this gene within non-target genotype.
